# The value of DNA methylation profiling in characterizing preeclampsia and intrauterine growth restriction

**DOI:** 10.1101/151290

**Authors:** Samantha L Wilson, Katherine Leavey, Brian Cox, Wendy P Robinson

## Abstract

Placental health is a key component to healthy pregnancy. Placental insufficiency (PI), inadequate nutrient delivery to the fetus, is associated with preeclampsia (PE), a maternal hypertensive disorder, and intrauterine growth restriction (IUGR), pathologically poor fetal growth. PI is more common in early-onset PE (EOPE) than late-onset PE (LOPE). However, the relationship between these disorders remains unclear. While DNA methylation (DNAm) alterations have been identified in PE and IUGR, these entities can overlap and few studies have analyzed these separately. This study aims to identify altered DNAm in EOPE, LOPE, and normotensive IUGR, validate these alterations, and use them to better understand the relationships between these related disorders.

Placental samples from a discovery cohort (43 controls, 22 EOPE, 18 LOPE, 11 IUGR) and validation cohort (15 controls, 22 EOPE, 11 LOPE) were evaluated using the Illumina HumanMethylation450 array. To minimize gestational age (GA) effects, EOPE samples were compared to pre-term controls (GA <37 weeks), while LOPE and IUGR were compared to term controls (GA >37 weeks). There were 1703 differentially methylated (DM) sites (FDR<0.05, Δβ>0.1) in EOPE, 5 in LOPE, and 0 in IUGR. Of the 1703 EOPE sites, 599 were validated in the second cohort. These sites cluster samples from both cohorts into 3 distinct methylation clusters. Interestingly, LOPE samples diagnosed between 34-36 weeks with co-occurring IUGR clustered with the EOPE methylation cluster. DNAm profiling may provide an independent tool to refine clinical diagnoses into subgroups with more uniform pathology. The challenges in reproducing genome-wide DNAm studies are also discussed.

## Introduction

Preeclampsia (PE) (OMIM 189800), a multi-system maternal hypertensive disorder of pregnancy, is the leading cause of maternal and perinatal morbidity and mortality worldwide, occurring in 2-8% of pregnancies (1). Intrauterine growth restriction (IUGR), defined as poor fetal growth due to an underlying pathology, often co-occurs with PE, but may also occur in normotensive pregnancies. Infants from pregnancies complicated by PE and/or IUGR are at risk for immediate and long-term adverse health outcomes (2,3). To date, there is no consistent test utilized to predict PE or IUGR prior to the onset of clinical symptoms. Protein biomarkers such as pregnancy-associated plasma protein A (PAPPA) and placental growth factor (PlGF) have been used to predict PE and/or IUGR (4); however, these methods may not be generalizable to other populations, as studies consist predominantly of high risk and Caucasian populations (5). Another limitation of current screening approaches is our poor understanding of the distinct pathological mechanisms and corresponding placental changes that may underlie these conditions.

Both PE and IUGR are heterogeneous in etiology, with many different factors contributing to these phenotypes (6,7,8). Risk factors for PE include genetic abnormalities, such as triploidy, some trisomies, and point mutations, as well as maternal health factors, such as obesity, pre-existing hypertension, and diabetes (9,10,11,12). Normotensive IUGR (nIUGR) can arise due to confined placental mosaicism, placental dysfunction, chronic inflammation of the placenta, poor maternal nutrition, smoking, stress, and other causes (12,13,14). Due to the heterogeneity in etiology of PE and IUGR, the ability to sub-classify placentas into more homogeneous groups can aid in our understanding of disease pathogenesis and prediction. For example, by defining ‘placental IUGR’ on the basis of a detailed scoring system for placental pathology, Benton *et al.* showed this subset was associated with very low maternal serum PlGF and also had the most severe perinatal and postnatal risks (15). In some cases, PE and nIUGR may represent two facets of a common underlying etiology, while in others the associated placental pathology and molecular changes may be distinct. Enforcing stringent criteria for defining and grouping samples may increase the reproducibility for reported molecular changes. For this study, we subdivide our samples into early-onset PE (EOPE), late-onset PE (LOPE), and nIUGR based on clinical obstetric criteria.

Molecular profiling has the potential to refine these clinically-defined group definitions further and aid in understanding the etiology of EOPE, LOPE, and nIUGR and their relationship to one another. Placental transcriptome profiling from pregnancies associated with PE and healthy controls provide evidence for multiple subtypes of PE (16). DNA methylation (DNAm) profiling is an alternative or complementary approach to gene expression profiling to identify subgroups of placental phenotypes. DNAm is more stable than mRNA and hence is less subject to changes with sample processing time (17); it may also retain a “memory” of earlier in utero exposures and hence be linked to early effects in the disease process.

We previously showed widespread DNAm alterations in EOPE (18) using the Illumina Infinium HumanMethylation450 Array (450K), measuring >480,000 CpG sites across the genome (19). We also demonstrated that placentas associated with confined placental trisomy 16, a condition that can be associated with PE, showed some overlapping changes with chromosomally normal EOPE, as well as a unique set of changes specific to the presence of the trisomy (20). Other groups have similarly found altered DNAm in PE and IUGR, though the differentially methylated sites or “hits” are not entirely consistent between studies (21,22,23,24,25,26,27). This inconsistency may be due to i) how sample groups are defined, ii) placental sampling differences, or iii) how groups define validated hits between the studies.

The aims of the present study were to build on our understanding of DNAm changes in placental insufficiency and to i) reanalyze our previous EOPE data with newer approaches; ii) extend our analysis to LOPE and nIUGR to investigate the potential DNAm relationships between the three pathologies using hierarchical clustering and, iii) validate differentially methylated sites in an independent cohort. We will also discuss challenges to validation and future directions for epigenetics in the placental biology field.

## Results

### Widespread DNAm changes are associated with EOPE but not LOPE and nIUGR in our Discovery cohort

Our first goal was to confirm our previous report of widespread changes in EOPE (18) and then to test for similar changes in LOPE and nIUGR using the same approach. We chose less stringent cutoffs for significance in this analysis (FDR<0.05 & Δβ>0.1) as compared to Blair *et al.* (2013) (FDR<0.01 & Δβ>0.125) (18) as our aim was to identify a larger number of differentially methylated sites that could be used for further validation. Based on these criteria, a total of 1703 sites were differentially methylated between EOPE and pre-term controls (**Figure 1**). As expected, the majority (261/286) of EOPE hits reported in Blair *et al.* were also identified as hits in this analysis. Differences between the two analyses are likely explained by the use of different normalization methods, correction for fetal sex in the present study, and the inclusion of a few additional samples in this study compared to the previous one.

**Figure 1.**
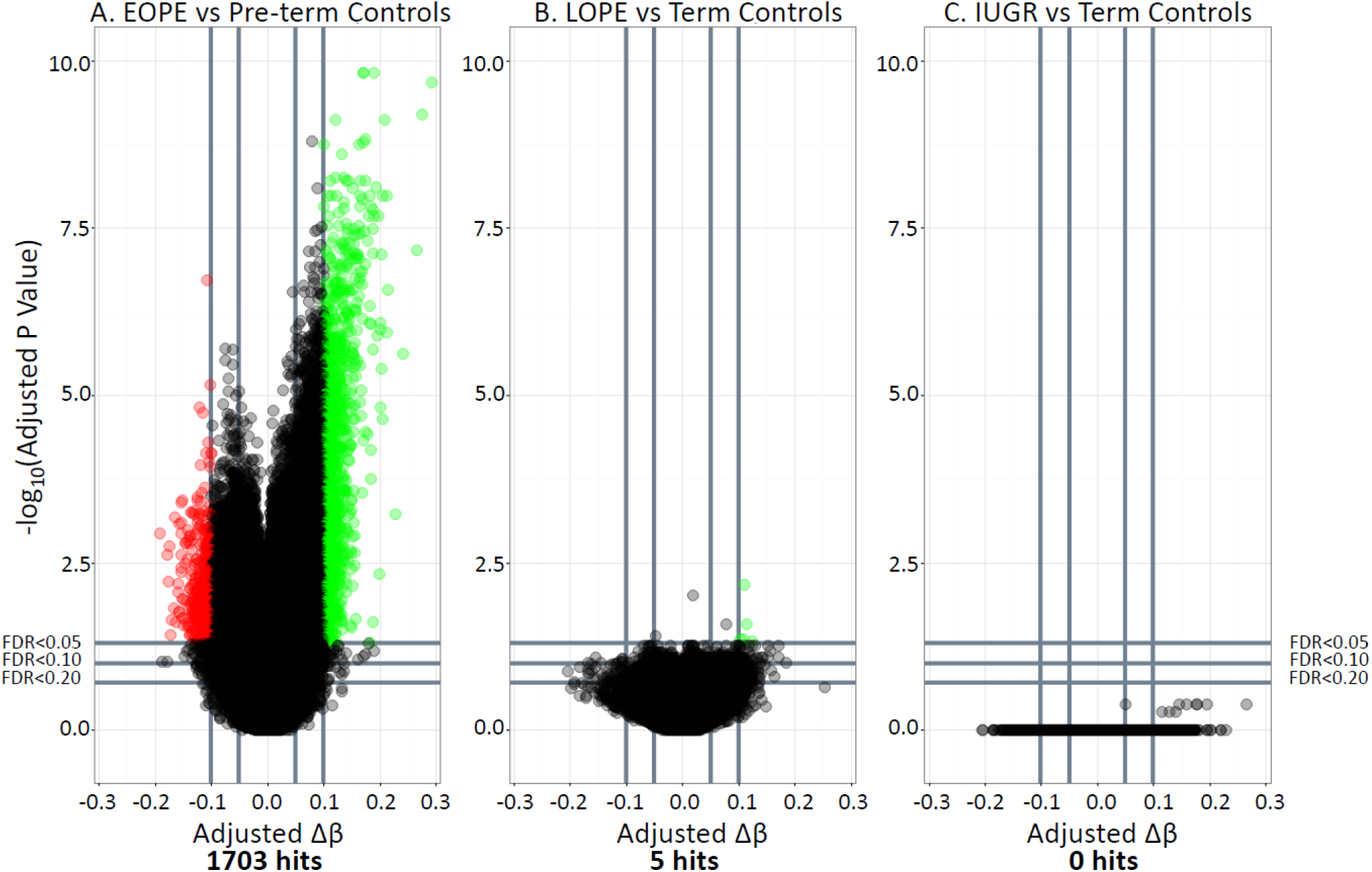
Volcano plots depicting differentially methylated sites between A) EOPE and pre-term controls, B) LOPE and term controls, and C) IUGR and term controls. −log10 of the adjusted p-value is plotted on the y axis and the change in DNAm (Δβ) is plotted on the x axis. Sites highlighted in red are hypomethylated in the pathology compared to controls. Sites highlighted in green are those that are hypermethylated in the pathology compared to controls.

We used the same approach to identify differential methylation associated with LOPE or nIUGR as compared to the healthy term control group. In contrast to the EOPE comparison, only 5 sites were differentially methylated between LOPE and term controls, and no sites were differentially methylated between nIUGR and term controls (**Figure 1**). The 5 differentially methylated sites between LOPE and term controls were not unique to LOPE, as they were also included amongst the 1703 sites identified as differentially methylated in EOPE. The weaker signal may be explained if only a few of the LOPE cases have an underlying pathology similar to that in the EOPE cases, driving these changes.

### Validation of the EOPE hits in an independent cohort

We next investigated if the EOPE hits from our discovery cohort could be validated in an independent cohort. We first tested whether the Δβ values in the discovery and validation cohorts were correlated using all sites that met an FDR<0.05 in the discovery cohort, without imposing an additional Δβ threshold. At these sites, the correlation was significant (R=0.62, p<2.2e-16, **Figure 2a**). This indicates that largely similar changes in DNAm are being observed in the EOPE placentas in both cohorts. Amongst the most highly significant hypermethylated sites (Δβ >0.15) in both cohorts were CpGs associated with *KRT15, FN1, TEAD3, JUNB, ST3GAL1, PKM2, NDRG1, PAPPA2, CHI3L2*, and *INHBA*. Many of these genes have previously been shown to have altered gene expression in preeclampsia and *JUNB* has been specifically implicated as a key player in the response to hypoxia in trophoblast cells (28). Amongst the most highly hypomethylated sites (Δβ <-0.10) in both cohorts were several sites associated with *FAM3B, SYNE1,* and *AGAP1*. However, it should be noted that there were also many sites with a high Δβ in the discovery cohort that had a much smaller or sometimes opposite direction Δβ in the validation cohort.

**Figure 2.**
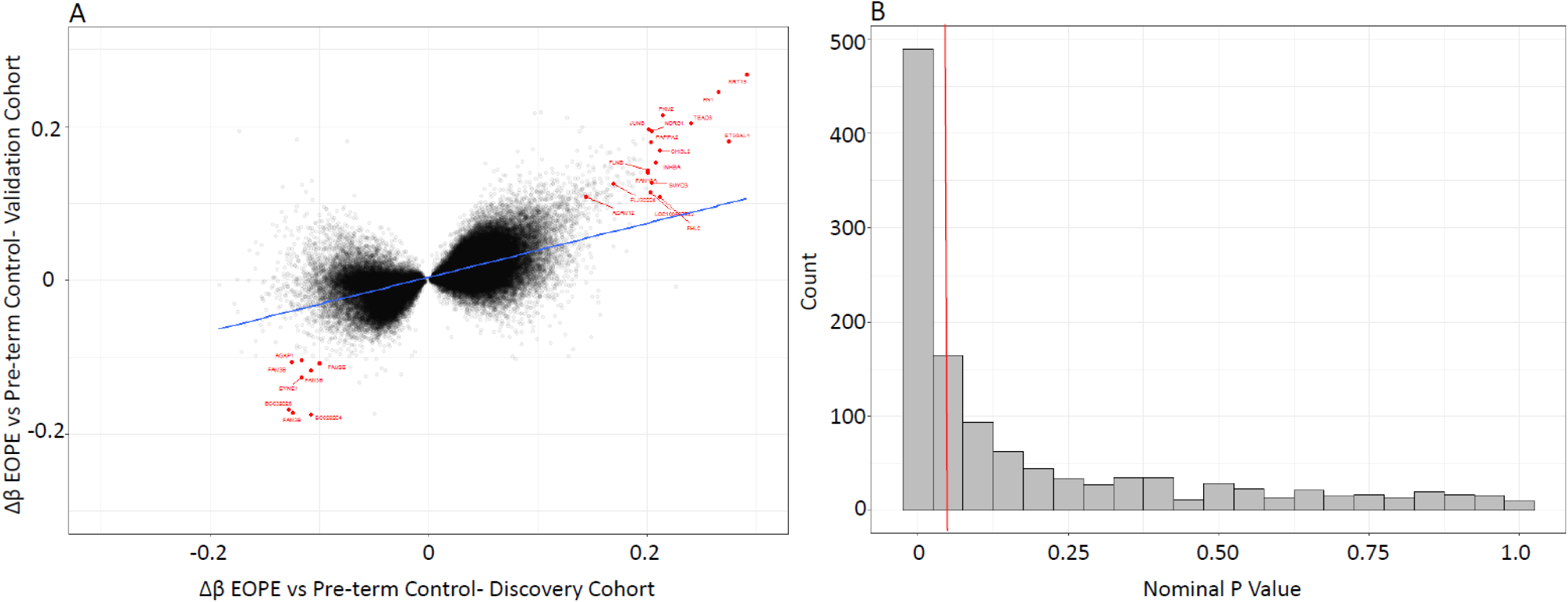
A) The correlation between the change in DNAm (Δβ values) between EOPE and pre-term controls, between the discovery and validation cohorts. The sites highlighted in red are the top sites labeled by the gene the CpG site is located in. B) P-value distribution of the 1703 EOPE hits from the discovery cohort, in the validation cohort.

To narrow down the original 1703 EOPE hits from the discovery cohort to a high-confidence hit list, we first asked, how many of these hits met similarly stringent criteria (FDR<0.05 and Δβ>0.1) in the validation cohort? We found that only 38 probes (2.2%) met these strict criteria. Using such arbitrary cutoffs in both populations and a strict definition for a “hit” may not be a powerful approach to assess the degree of overlap in the data. Furthermore, requiring assay-wide correction for multiple testing in the validation cohort is overly conservative and reduces power. Running the linear regression on only the 1703 sites differentially methylated in the discovery cohort reduces the number of multiple test corrections needed in the validation cohort. Based on the distribution of nominal p-values among the 1703 EOPE associated sites in the validation cohort, shown in **Figure 2b**, there are many more sites that meet a nominal p-value<0.05 than expected, even if these do not meet a multiple test correction. Hence, we opted to use a nominal p-value<0.05 and a change in DNAm in the same direction as the discovery cohort to define validated (i.e. high confidence) hits. Based on these criteria, 599 of the 1703 (35.1%) EOPE hits were considered to be validated (**Figure 2b**). This is higher than what we would expect by chance (p=0.0001). This reproducibility rate was similar to the rate reported by Yeung *et al.* (2016), who validated their own differentially methylated regions with our published cohort (Blair *et al.* (2013))(29). As PE and IUGR are heterogeneous conditions, it is possible that the reproducibility rate may be affected by the samples chosen for each cohort. We were interested in whether the samples in the cohorts were similarly correlated (i.e. is one cohort more heterogeneous than the other). We investigated these correlations in the control samples (Term and Pre-term) (**Supplementary Figure 2**) and the EOPE samples (**Supplementary Figure 3**). In both pathologies, samples in the discovery cohort were more heterogeneous than the samples in the validation cohort.

These validated sites were not enriched for any gene ontology terms using ermineJ, with a 450K array specific background (30). A list of these sites and relevant gene information can be found in **Supplementary Table 1**. These sites include ones associated with genes known to be relevant to EOPE from gene expression studies including *CGA*, *INHBA*, *PAPPA2*, and *ADAM12*.

### Effects of varying the validation criteria

To evaluate the effect of varying FDR and Δβ cutoffs to establish the most ‘reproducible’ results, we plotted the percentage of probes that showed Δβ concordance in directionality between the validation cohort and the discovery cohort using different FDRs and Δβ thresholds in the discovery set (**Supplementary Figure 1a**). Different FDR thresholds did not influence DNAm concordance rate when the Δβ thresholds were above 0.2. FDR thresholds appear to be more important when trying to identify small changes in DNAm. We also investigated the number of hits that each threshold would obtain. **Supplementary Figure 1b** plots the number of hits at each FDR and Δβ cutoff. Allowing smaller changes in DNAm produces many more hits, but with a lower reproducibility rate. This is likely because this is in the range of normal variability for a site. This highlights the importance of considering both the biological and statistical thresholds depending on the magnitude of the anticipated DNAm change and the overall research objective.

### Hierarchical Clustering

Next, we evaluated the degree to which the 599 validated sites can discriminate EOPE from all other placentas in both cohorts, using hierarchical clustering (**Figure 4**). Although we expected an EOPE methylation cluster to be defined, since we are clustering based on our EOPE hits, this approach can tell us about the relationships between individual samples and, furthermore, allow comparisons to term controls, and LOPE and nIUGR cases which were not involved in the selection of these EOPE hits. Interestingly, both cohorts clustered into 3 stable methylation clusters, rather than just two as we had expected (**Figure 3**). When the cohorts were clustered on the 599 validated hits separately, methylation cluster 1 included almost all EOPE suggesting a consistent phenotype in this group. In the discovery cohort cluster, 6 LOPE samples clustered with the EOPE samples. In the validation cohort, 7 LOPE samples and 1 pre-term control clustered with the EOPE samples. Additionally, in both cohorts, methylation sub-clusters were identified within the larger EOPE group (methylation cluster 1), which were also stable and significantly different from one another, suggesting a possible further subdivision or distinct groups within methylation cluster 1. There was no obvious difference between these subclusters clinically (including sex, ethnicity, disease severity etc.); however, gestational age was decreased in the EOPE methylation subcluster 1 compared to subcluster 2 (p<0.01) in the validation cohort (**Table 1**).

**Figure 3.**
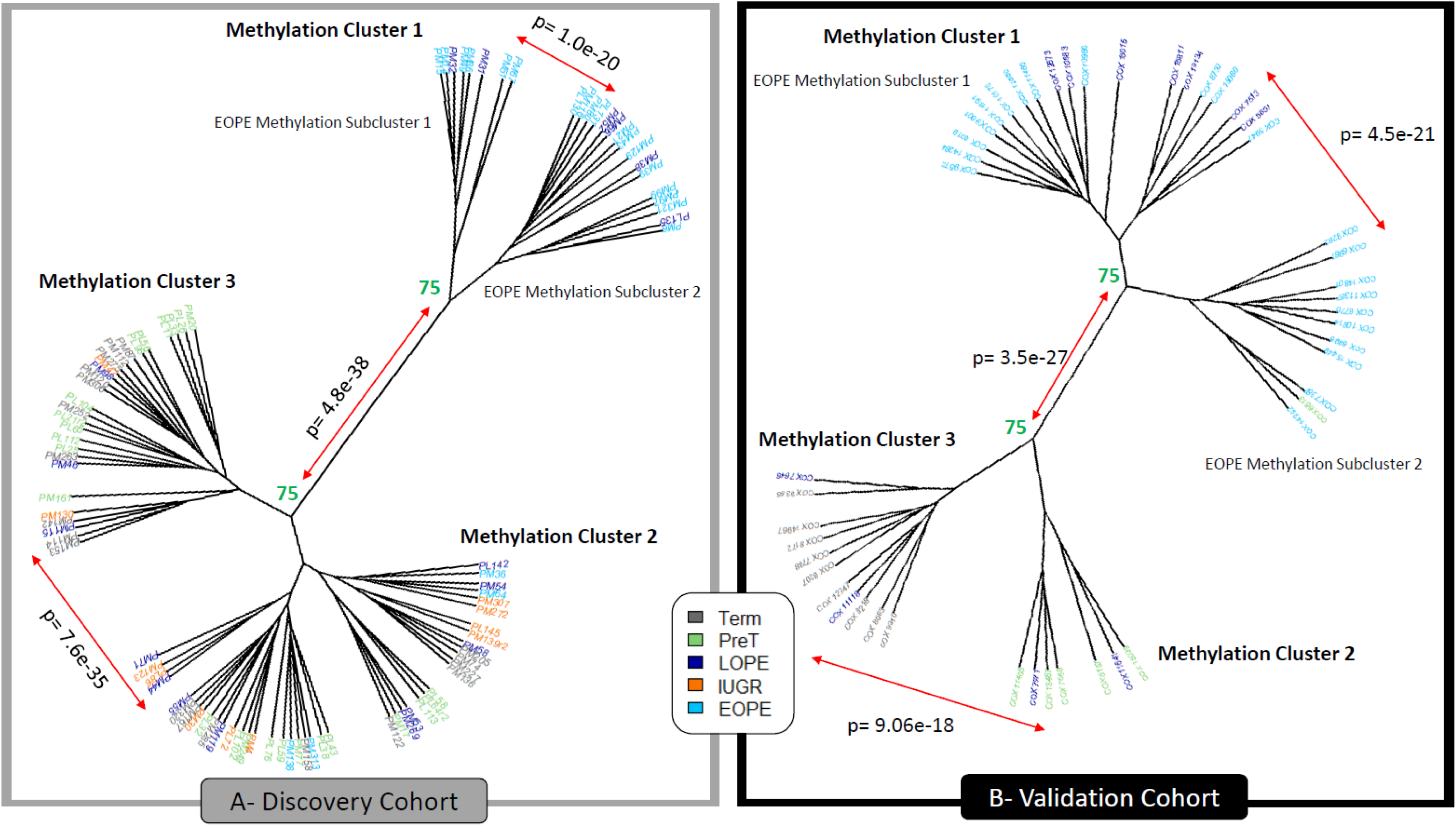
Hierarchical clustering (Euclidean) on the 599 validated hits in both the discovery (left) and validation (right) cohorts. Numbers represent the percentage of times these clusters formed when using 1000 iterations with pvclust. Those highlighted in green are considered stable, where clusters formed >75% of the time. Those highlighted in red were unstable. p values signify clusters are significantly different from one another.

**Table 1.** Clinical information on samples assigned to methylation cluster 2 compared to methylation cluster 3 and samples assigned to EOPE methylation subcluster 1 and EOPE methylation subcluster 2.

| DISCOVERY COHORT |  |  |  |  |
| --- | --- | --- | --- | --- |
|  | Methylation Cluster 2 | Methylation Cluster 3 | EOPE Methylation Subcluster 1 | EOPE Methylation Subcluster 2 |
| N= | 43 | 27 | 8 | 16 |
| IUGR Status (IUGR/Total) | 10/43 (23%)* | 2/27 (7%) | 7/8 (88%) | 14/16 (88%) |
| Fetal Sex (F/Total) | 22/43 (51%) | 11/27 (41%) | 5/8 (63%) | 3/16 (19%)* |
| Gestational Age (weeks) (range (mean)) | 25.0-40.0 (36.0) | 28.0-41.3 (35.7) | 24.9-36.0 (31.7) | 26.0-37.3 (32.6) |
| Fetal Birth Weight (SD) (range (mean)) | -2.78 – 3.77 (-0.44)** | -2.16 – 1.70 (0.18) | -2.90 - -1.17 (-1.87) | -8.19 - -0.58 (-2.26) |
| Chronic Hypertension (CH/Total) | 7/43 (16%) | 2/27 (7%) | 2/8 (25%) | 5/16 (31%) |
| Diabetes (Pre-existing or Gestational) (Diabetes/Total) | 2/43 (5%) | 0/27 (0%) | 1/8 (13%) | 1/16 (6%) |
| Chorioamnionitis (CA/Total) | 7/43 (16%) | 5/27 (19%) | 0/8 (0%) | 0/16 (0%) |
| Premature Rupture of Membranes (PPROM/Total) | 3/43 (7%) | 4/27 (15%) | 0/8 (0%) | 0/16 (0%) |
| Ultrasound Findings (Findings/Total) | 7/43 (16%) | 2/27 (7%) | 3/8 (38%) | 4/16 (25%) |
| Placental Pathology Noted (Notes/Total) | 17/43 (40%) | 7/27 (26%) | 4/8 (50%) | 10/16 (63%) |
| VALIDATION COHORT |  |  |  |  |
|  | Methylation Cluster 2 | Methylation Cluster 3 | EOPE Methylation Subcluster 1 | EOPE Methylation Subcluster 2 |
| N= | 7 | 11 | 19 | 11 |
| IUGR Status (IUGR/Total) | 0/7 (0%) | 0/11 (0%) | 11/19 (58%) | 7/11 (64%) |
| Fetal Sex (F/Total) | 1/7 (14%) | 5/11 (45%) | 9/19 (47%) | 7/11 (64%) |
| Gestational Age(weeks) (range (mean)) | 30.0-37.0 (33.0)*** | 37.0-40.0 (38.4) | 27.0-37.0 (32.8)*** | 26.0-34.0 (29.8) |
| Fetal Birth Weight (SD) (range (mean)) | -0.48 – 0.50 (-0.06) | -1.11 – 3.60 (0.34) | -2.27 – 2.92 (-1.12) | -2.46 - -0.86 (-1.59) |
| Chronic Hypertension (CH/Total) | 1/7 (14%) | 1/11 (9%) | 7/19 (37%) | 3/11 (27%) |

\* p<0.1, \*\* p<0.05, \*\*\*p<0.01

The remaining non-EOPE samples also clustered into two methylation clusters in both cohorts. We refer to these clusters as methylation clusters 2 and 3. Methylation cluster 3 in both cohorts was predominantly composed of controls. Within the discovery cohort, methylation cluster 2 consisted of the majority of the nIUGR and LOPE cases, a few EOPE cases, and some pre-term and term controls; Decreased birthweight (p<0.01) and a trend towards increased IUGR diagnosis (p<0.1) was observed in methylation cluster 2 vs. 3. In the validation cohort, methylation cluster 2 consisted of pre-term controls and LOPE samples and was associated with decreased gestational age (p<0.01) (**Table 1**).

### Cluster Gene ontology

As it was unexpected that the control samples divided into two distinct methylation clusters, and the EOPE samples divided into two distinct subclusters, we were interested in investigating the differences between the two control methylation clusters (methylation clusters 2 and 3) and between the EOPE methylation subclusters (EOPE methylation subclusters 1 and 2). Linear regression was used on the 599 persistent hits, accounting for fetal sex, to identify DNAm differences between methylation cluster 3 and methylation cluster 2 and between EOPE methylation subcluster 1 and EOPE methylation subcluster 2. Using a nominal p-value<0.05, 244 of those sites were differentially methylated between methylation cluster 3 and methylation cluster 2 in both cohorts. Information on these sites can be found in **Supplementary Table 1**. There was no enrichment in gene ontology terms in either ermineJ (with a 450K array specific background), or DAVID. Between EOPE methylation subcluster 1 and EOPE methylation subcluster 2, 207 sites were differentially methylated in both cohorts. Information on these sites can be found in **Supplementary Table 2**. There was no gene ontology enrichment by ermineJ and symporter activity was the only gene ontology term in DAVID to meet multiple test corrections.

## Discussion

We previously reported widespread changes in DNAm associated with EOPE (18). In the present study, we extend this analysis to LOPE and nIUGR; however, using the same approach, we were unable to identify DNAm changes that were unique to these groups. While these latter comparisons were limited by small sample size, we were able to obtain significant associations with EOPE with similarly small sample sizes. The reduced number of changes in the LOPE and nIUGR groups can occur for two main reasons: 1) there may be much more limited placental pathology with these diagnoses and the phenotype is largely driven by maternal factors, or 2) they may be more heterogeneous etiology thereby limiting power to detect changes in the group as a whole. If we want to improve biomarker discovery in these groups, we may need to identify more homogeneous subgroups using a combination of clinical parameters, pathology reports and/or biomarkers themselves, along with larger sample sizes.

The LOPE samples in the discovery cohort that clustered with the EOPE samples all presented with PE between 34.0 weeks and 35.9 weeks gestation and had co-occurring IUGR. While it’s possible that PE symptoms were present but not diagnosed until after 34.0 weeks, there may also be inaccuracies in dating the pregnancy and/or there is simply a grey zone in the distinction between EOPE (placenta-driven) and LOPE (maternal-health driven). There were also four cases of EOPE within the discovery cohort that did not cluster with other EOPE cases. One was diagnosed with hemolysis elevated liver enzymes and low platelet (HELLP) syndrome and delivered at 33.3 weeks gestation, one also had chorioamnionitis (which may have contributed to early delivery); one had preexisting hypertension and was diagnosed early but did not deliver until 37 weeks and hence may have been milder in presentation; the fourth was delivered at 33.3 weeks with no other placenta or maternal health notes. Of note, none of the EOPE cases that clustered outside of the EOPE cluster had co-occurring IUGR and were generally diagnosed at close to 34 weeks gestation. Thus, the presence of IUGR in cases diagnosed between 32 and 36 weeks may be the more defining feature as to whether an altered placental DNAm profile is observed or not. Powers *et al.* (2012) showed that there are two types of PE pregnancies: those with and without altered angiogenic factors (32). As alterations in the angiogenic factors have also been observed in IUGR cases (33,34), Myatt and Roberts suggested that an imbalance in these factors may represent a measure of placenta growth, development, and function (35). As such, the EOPE cases clustering outside the EOPE methylation cluster may be more likely related to other contributing factors than placental dysfunction.

While we expected to see an EOPE methylation cluster, as the validated hits were chosen based on differentially methylated sites between EOPE and pre-term controls, we were surprised that both the EOPE and control groups each formed two subclusters. The driving differences between these subclusters were not clear, though the tendency to lower gestational ages and fetal birth weights (SD) in cluster 2 could suggest features linked to preterm birth (**Table 1**). The presence of two distinct subclusters within the EOPE methylation cluster could reflect PE severity, or perhaps unmeasured factors such as medical treatments given or duration of hypertension. Unfortunately, we had insufficient information on the treatment of each case to evaluate the influence of medical care on the placental methylation profile.

While altered placental DNAm has been reported for pregnancies complicated by PE (21,22,23,25) and IUGR (24,26,27), only one study validated their findings, using the same technology, in an independent cohort (29). In this study, 35.1% (N=599) sites were found to be differentially methylated between EOPE and pre-term controls in both the discovery and validation cohorts using validation criteria of a nominal p<0.05 and change in DNAm in the same direction as the discovery cohort. The extent of validation, however, is dependent on the initial criteria chosen to define ‘hits’, the criteria for validation, the similarity of the populations of samples, the size of the study populations (power to detect changes), and the similarity in processing the samples. In our study, the validation cohort was from a roughly similar urban population (Vancouver vs. Toronto) from the same country (Canada). We also tried to minimize technical factors that may influence results by using similar placental sampling protocols, processing the arrays with a subset of the discovery and validation cohorts on the same microarray chips at the same time, with the same technicians, and using the same pre-processing methods on the raw data. Even with these considerations, a significant number of our original hits were not validated. This may be because of chance variation in causes of PE and IUGR in the two cohorts due to limited sample size. Additionally, there are genetic, environmental, and maternal factors that pre-dispose a pregnancy to developing placental insufficiency, which may have varied between populations.

Changes in DNAm could mean i) an average change in DNAm across our sample, or ii) a change in the cell type proportions within a sample, as DNAm varies widely across different cell types (40). In the context of EOPE, DNAm alteration may reflect a combination of altered gene expression pathways associated with PE (ex. related to known effects such as oxidative stress and altered angiogenesis (7,32)) or altered cell type proportions related to PE pathology (ex. decreased proliferation of extravillous trophoblast cells or alterations to the rate of trophoblast proliferation (41)). As cell-type specific profiles have not been developed for all placental cell types, it is not possible to use the DNAm profile to estimate cell proportions, as it has been applied to blood (42). While reference-free methods for deconvolution of cell proportions have been developed (43), these methods remove variance within the data attributed to cell composition but cannot inform us of what cell types specifically are altered in EOPE.

### Summary

Our data demonstrate some of the challenges in identifying changes specific to clinically defined etiologies. Heterogeneity and milder phenotypes in LOPE and nIUGR likely limit the power to detect differences using a differential methylation type approach and mask the subset of cases that do exhibit altered pathology (based on sample clustering using our EOPE defined hits). An alternative approach may be to reduce the dimensions in the data by 1) removing non-variable probes across all cell types (36), 2) focusing on alterations in pathway modules, as in weighted gene co-expression network analysis (37), or 3) evaluating differentially methylated regions (DMRs) rather than individual CpG sites to combat multiple test correction (38,39).In contrast, the more severe pathology underlying EOPE results in many readily detected DNAm changes. However, even in the case of EOPE, where many large changes in DNAm are identified and can be validated based on a nominal p-value<0.05 in an independent cohort, those sites selected for having the highest magnitude of change rarely showed the same degree of difference in the second cohort. Techniques to reduce the dimensions in the data should be developed and utilized, focusing on altered pathways instead of specific changes, which may help in identifying subtypes of PE and IUGR to guide and change management in a useful way.

In conclusion, whether in the context of PE, or other heterogeneous diseases, DNAm may be a useful tool to independently and qualitatively classify pathological groups prior to analysis. This method may aid in creating more robust prediction algorithms for predicting pathology versus controls. Further studies with larger sample sizes and additional clinical variables are needed to confirm the presence of multiple subtypes of placental-mediated PE, and what is driving these different subtypes.

## Methods

### Sample Information

#### Placental Sampling

Chorionic villi samples were obtained from 2-3 sites in the placenta, each from distinct cotyledons, as previously described (18). Infarcts or necrotic regions of the placenta were avoided in sampling and DNA was extracted from each site, using a standard salting out method and pooled together in equal amounts to give a more accurate representation of the placenta’s molecular profile. The NanoDrop 1000 spectrophotometer (ThermoScientific, Wilington,DE,USA) was used to assess DNA purity and concentration.

#### Discovery Cohort

The discovery (Vancouver) cohort consisted of 22 EOPE, 18 LOPE, 11 nIUGR and 43 control placentas (**Table 2**). Ethics approval from both the University of British Columbia and BC Women’s and Children’s Hospital ethics committees in Vancouver, BC, Canada, was obtained (H04-704488). Placental samples were obtained with consent from patients from the Medical Genetics as well as the Obstetrics and Gynecology departments. Clinical information, including gestational age at delivery, fetal sex, fetal birth weight, and maternal age were collected. Criteria for exclusion were multi-fetal pregnancies and fetal and/or placental chromosomal abnormalities. A subset of 18 EOPE samples and 19 pre-term control samples in this study was previously used in Blair *et al.* (2013)(18).

**Table 2.** Discovery and validation cohort clinical information

| Discovery Cohort |  |  |  |  |  |
| --- | --- | --- | --- | --- | --- |
|  | EOPE<br>(mean) | LOPE<br>(mean) | IUGR<br>(mean) | Pre-term "Control"<br>(mean) | Term Control<br>(mean) |
| N= | 22 | 18 | 11 | 24 | 19 |
| Fetal birth weight- Standard deviation range | -8.18 – 3.77<br>(-1.65)<sup>1</sup> | -2.9 – 2.57<br>(-0.96)<sup>2</sup> | -2.57 – -1.22<br>(-1.99)<sup>3</sup> | -1.61 – 3.23<br>(0.51) | -0.94- 0.98<br>(-0.09) |
| Fetal Sex (M:F) | 14:8 | 10:8 | 4:7 | 16:8 | 9:10 |
| Maternal Age (years) | 19.7-42.9<br>(33.3) | 23.1-41.3<br>(34.0) | 33.3-38.0<br>(34.3) | 22.2-41.1<br>(32.5) | 30.0-40.2<br>(34.9) |
| Gestational Age (weeks) | 24.9-38.4<br>(32.0) | 34.6-41.3<br>(37.4) | 34.6-38.0<br>(36.6)<sup>4</sup> | 25.0-36.7<br>(32.6) | 37.3-39.9<br>(38.4) |
| Validation Cohort |  |  |  |  |  |
|  | EOPE<br>(mean) | LOPE<br>(mean) | IUGR<br>(mean) | Pre-term "Control"<br>(mean) | Term Control<br>(mean) |
| N= | 22 | 11 | 0 | 6 | 9 |
| Fetal birth weight- Standard deviation range | -2.46- 0.01<br>(-1.40)<sup>5</sup> | -2.92- 0.01<br>(-0.84)<sup>6</sup> | 0 | -1.02- 0.05<br>(-0.16)<sup>7</sup> | -1.11- 3.60<br>(0.61) |
| Fetal Sex (M:F) | 9:13 | 7:4 | 0 | 4:2 | 6:3 |
| Gestational Age (weeks) | 26.0-34.0<br>(30.5) | 35.0-37.0<br>(36.4)<sup>8</sup> | 0 | 27.0-33.0<br>(30.8) | 38.0-40.0<br>(38.8) |

Discovery Cohort: <sup>1</sup> p= 6.9e-6 vs PTB "Control", <sup>2</sup> p=0.014 vs Term Control, <sup>3</sup> p= 1.7e-5 vs Term Control, <sup>4</sup> p=3.3e-4 vs Term Control Secondary Cohort: <sup>5</sup> p=0.004 vs PTB "Control", <sup>6</sup> p=0.0005 vs Term Control, <sup>8</sup> p=0.0001 vs Term Control Between Cohorts: <sup>7</sup> p=0.04 PTB (discovery) vs PTB (secondary)

PE was defined according to the Society of Obstetricians and Gynecologists of Canada (SOGC) criteria as one of i) hypertension (BP>140/90mm Hg) and proteinuria (>300mg/day) arising after 20 weeks gestation (44); ii) HELLP syndrome without hypertension or proteinuria; or iii) eclamptic seizure without previous hypertension or proteinuria. Preeclampsia was separated into early and late onset given the clinical evidence that these may be associated with distinct risk factors and outcomes (6). Early-onset preeclampsia (EOPE) was defined as a diagnosis of PE prior to 34 weeks gestation, while LOPE was PE diagnosed after 34 weeks (6). nIUGR was defined as fetal birth weight <3^rd^ percentile accounting for both fetal sex and gestational age at delivery or fetal birth weight <10^th^ percentile accounting for both fetal sex and gestational age at delivery, with additional findings for poor fetal growth (44). As birthweight is strongly correlated with gestational age, we use the standard deviation of the birth weight corrected for fetal sex and gestational age (45).

Technical batch effects, related to the plate, microarray chip, and sample position on the Illumina chip are potential confounding factors within our data. Our samples were run in various batches over a 4 year period, and pathology and gestational age were partially confounded with batch as EOPE and preterm controls were largely run earlier. In this situation, correction for batch effects can introduce spurious findings (31) (**Supplementary Figure 4 and Figure 5**). We, therefore, instead compared EOPE to pre-term birth controls and LOPE/nIUGR to term controls only, which were relatively matched for batch, and thus the confounding by GA and its interaction with batch was minimized. We acknowledge that some of the differentially methylated sites that we found may be due to technical artifacts, but focusing on those hits that are reproduced in the validation cohort largely eliminated these effects.

As placental DNAm changes with gestational age, the comparison groups included placentas from healthy term births (>=37 weeks) and pre-term births (<37wks) with normally grown babies and no evidence of maternal hypertension. EOPE placentas were compared to 24 pre-term controls (as in Blair *et al. 2013*). LOPE and nIUGR placentas were compared to a separate set of 19 term control placentas. This was to test for overlap between DNAm changes identified for LOPE and nIUGR with those for EOPE, we did not want the use of a control group driving any potential overlap. To reduce the chance of differences being driven by the preterm birth group, we used placentas from pre-term births from a variety of etiologies (ex. premature rupture of the membranes, incompetent cervix, chorioamnionitis), while any term control samples with evidence of pathology involving the chorionic villi were excluded.

#### Validation Cohort

The validation (Toronto) cohort consisted of 22 EOPE, 11 LOPE, and 15 control placentas (**Table 2)**. For the validation cohort, placental samples were purchased through the Research Centre for Women’s and Infants’ Health BioBank (Mount Sinai Hospital), details in the sample processing can be found in Leavey *et al* (2016) (46). DNA was extracted from the pooled placental tissue by ethanol precipitation using the Wizzard^®^ Genomic DNA Purification Kit (Promega). Gestational age at delivery and fetal birth weight were collected for each case. For this cohort, ethics approval was obtained from both Mount Sinai Hospital (#13-0211-E) and the University of Toronto (#29435).

The validation cohort represents a subset of samples from the Leavey *et al.* (2016) study (46). PE was defined as BP > 140/90mm Hg after 20 weeks gestation and proteinuria >300mg/day or >2+ by dipstick (47). As the time of diagnosis was unknown, we subdivided the PE samples from this cohort into EOPE and LOPE based on the gestational age at delivery. Exclusion criteria included diabetes, sickle cell anemia, morbid obesity, and multi-fetal pregnancies. The division of the term and pre-term controls was also done in the validation cohort, which consisted of 6 pre-term control and 9 term control placentas (**Table 2**).

### DNA methylation Analysis

750ng of DNA purified using the Qiagen blood and tissue kit was bisulfite converted using the EZ DNA Methylation Kit (Zymo Research, Irvine, USA). Samples were run on the Illumina HumanMethylation450 BeadChip array platform (450K), measuring DNAm at 485,512 CpG sites across the genome (19). The samples, run on 27 chips in 4 batches, were hybridized to the microarray chip as per the manufacturer’s protocol, and microarray chips were scanned by a HiScan 2000 (Illumina). To minimize any effects of sample processing, arrays were run in the same batch and with the same operators as a subset of the samples from the discovery cohort (Chips 5013, 5015, 3024,3037,3038,3110, **See Supplementary Figure 4 and Figure 5)**. This DNA methylation data for the discovery and validation cohorts is available from the Gene Expression Omnibus (GEO) database under the accession numbers [GEO### In Process] and GSE98224, respectively.

Raw data (IDAT Files) were read into R statistical software, version 3.2.4, where functional normalization (48), background subtraction, and colour correction were performed. Blair *et al.* (2013), previously used subset within-array normalization (SWAN). Functional normalization performs all the benefits of SWAN normalization and, in addition, utilizes the 848 control probes on the array to mediate changes in DNAm that are due to technical effects (48). Bad quality probes and those that had a missing beta value in > 5% of samples or a detection p-value<0.01 were removed from the analysis (Discovery N=1,402, Validation N=1,115). To minimize fetal sex effects, probes on the X and Y chromosomes (Discovery N= 11,648, Validation N=11,302), as well as probes that cross-hybridize to the X and Y chromosomes (Discovery N= 11,412, Validation N=10,734), and probes containing a SNP at the CpG of interest were also removed (Discovery N= 19,957, Validation N= 20,398) (49). This left 440,093 CpG sites for analysis in the discovery cohort and 441,963 CpG sites in the validation cohort (**Supplementary Table 4**).

### Differential Methylation Analysis

All statistical analyses were performed using R version 3.2.4. Differentially methylated sites were identified using statistical, i.e. false discovery rate (FDR) <0.05, and biological, i.e. a change in DNAm (Δβ)>0.1, criteria. We corrected for fetal sex in our linear regression model, but we did not adjust for fetal birth weight, as it is closely related to pathology. As our groups were matched to controls of a similar gestational age, our final model where DNAm alterations were identified took into account fetal sex only. Only those sites that met both these criteria were then evaluated in the validation cohort. In this case, linear regression was used, and sites were considered to be persistent hits if the nominal p-value <0.05 and the change in DNAm was in the same direction as the discovery cohort. Bonferroni correction p <0.05 was also used to investigate how many hits would be validated with a more stringent threshold. To compare the heterogeneity of the samples in each cohort, sample by sample Pearson**’**s correlations were performed, and the average correlation of each sample was compared between the two cohorts by Student**’**s t-test separately for the control (Term + Pre-term) and EOPE samples.

To investigate whether the 42 Bonferroni corrected hits and the 599 nominal p-value validated hits were more than would be expected by chance, 1703 sites (number of EOPE hits in the discovery cohort) were randomly sampled from the validation cohort data and run through a linear model, correcting for fetal sex. One thousand permutations were run and the number of sites that met a nominal p-value<0.05 in each iteration was recorded. The number of randomly sampled sites to meet a nominal p-value<0.05 were compared to the actual number of sites that validated in our data (N=42 (Bonferroni corrected) and N=599 (nominal p-value)).

### Clustering Analysis

Hierarchical clustering was performed on the persistent hits to investigate whether samples clustered according to their clinically diagnosed pathology, or whether DNAm profiling could suggest an improved definition of pathological groups. The pvClust package in R (50) assessed how stable any resulting clusters were, using 1000 iterations. The sigClust2 package (51) determined if any clusters were significantly different from one another, also using 1000 iterations. To investigate whether differences in DNAm between the clusters were enriched for any specific pathway(s), linear regression was used to identify differentially methylated sites between clusters. Differentially methylated sites were annotated to genes using the Price *et al.* Annotation, closest transcriptional start site (49), and then inputted into ermineJ, a gene ontology tool (30). ErmineJ allows us to input a background gene list specific to the Illumina 450K array, accounts for multifunctionality (gene ontology terms that appear frequently due to the number of genes involved in the pathway), and allows for multiple iterations to be run to strengthen the power of the analysis.

## Acknowledgements

We would like to thank Kristal Louie, Dr. J Schuetz, Dr. S Langlois, and Dr. P von Dadelszen for recruiting patients for this study; Ruby Jiang for doing the placental dissections and DNA extractions, and Dr. M Penaherrera with help in both patient recruitment, and assistance in running the 450K arrays. Thanks to all members of the Robinson lab for reviewing and providing valuable feedback on the manuscript.

---

**Supplementary Figure 1.**
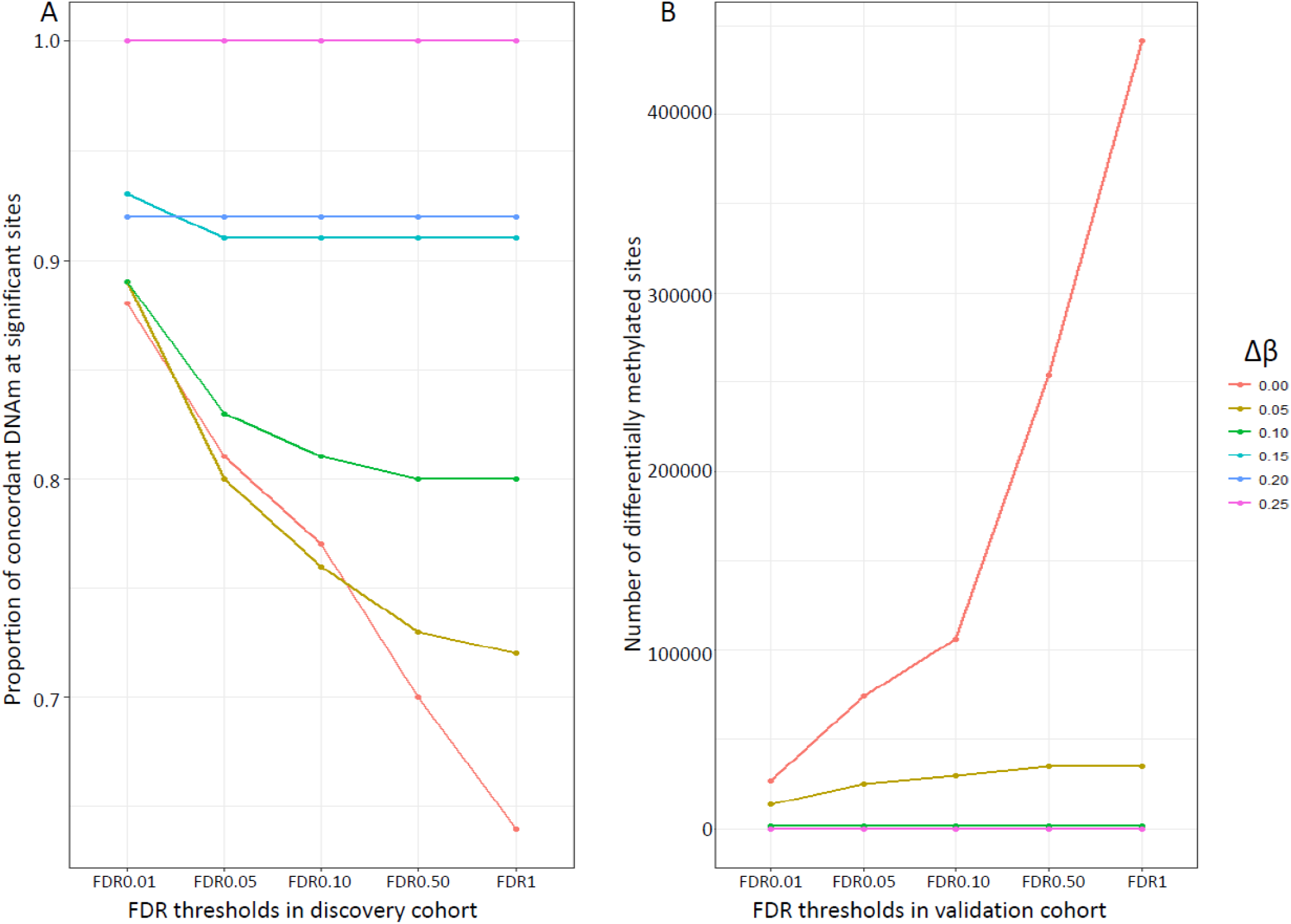
A) The concordance rate (sites where DNAm change is in the same direction in both cohorts) for different FDR and Δβ thresholds. Concordance rates (%) are plotted on the y-axis and FDR thresholds are on the x-axis. B) The number of identified hits for different FDR and Δβ thresholds. The number of hits is plotted on the y-axis and the FDR thresholds are on the x-axis.

**Supplementary Figure 2.**
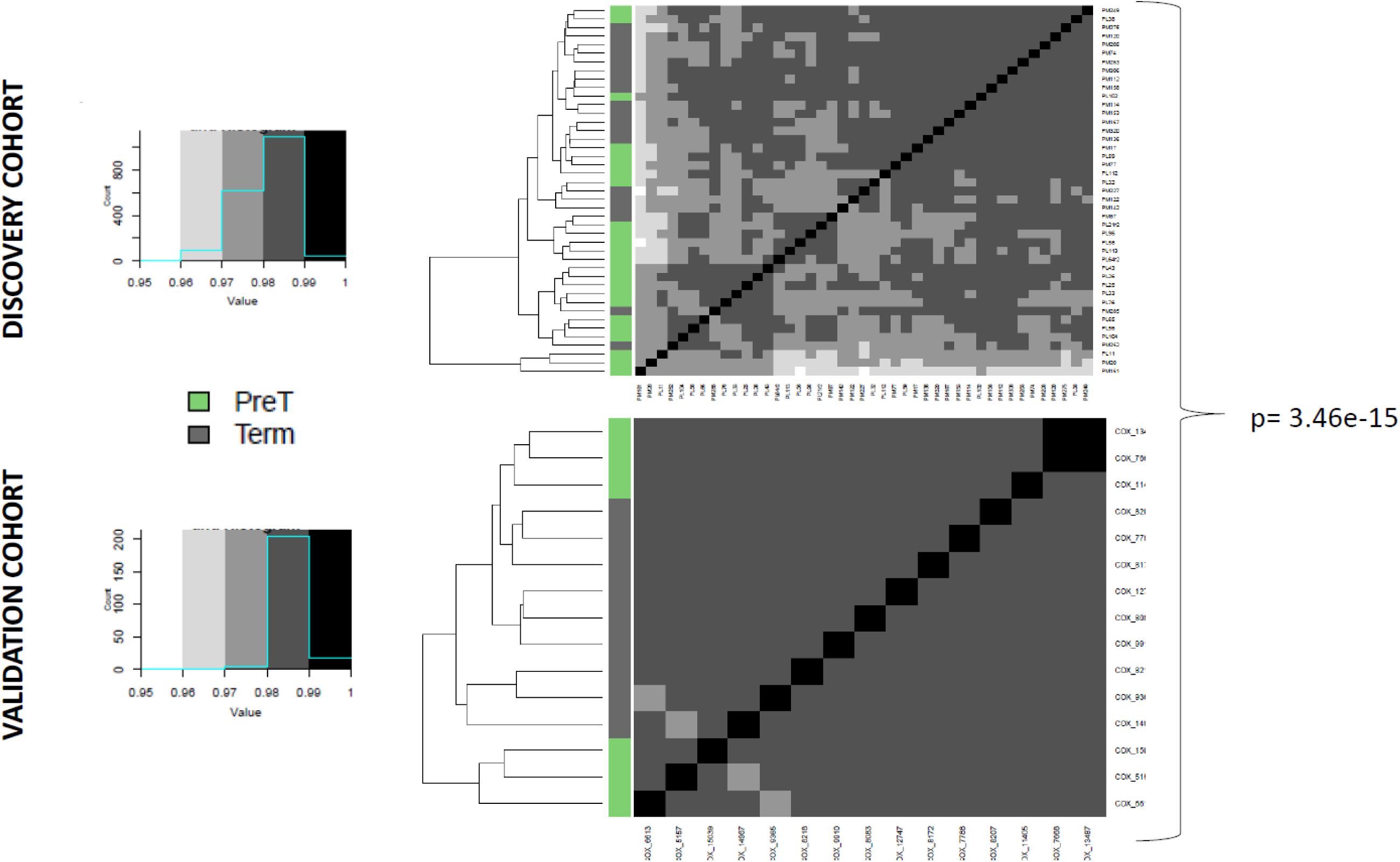
Sample-sample correlations for both the discovery and validation cohort in the control samples (both term and pre-term). P-value from Student’s t-test indicates that the discovery cohort is more heterogeneous (less correlated) than the validation cohort.

**Supplementary Figure 3.**
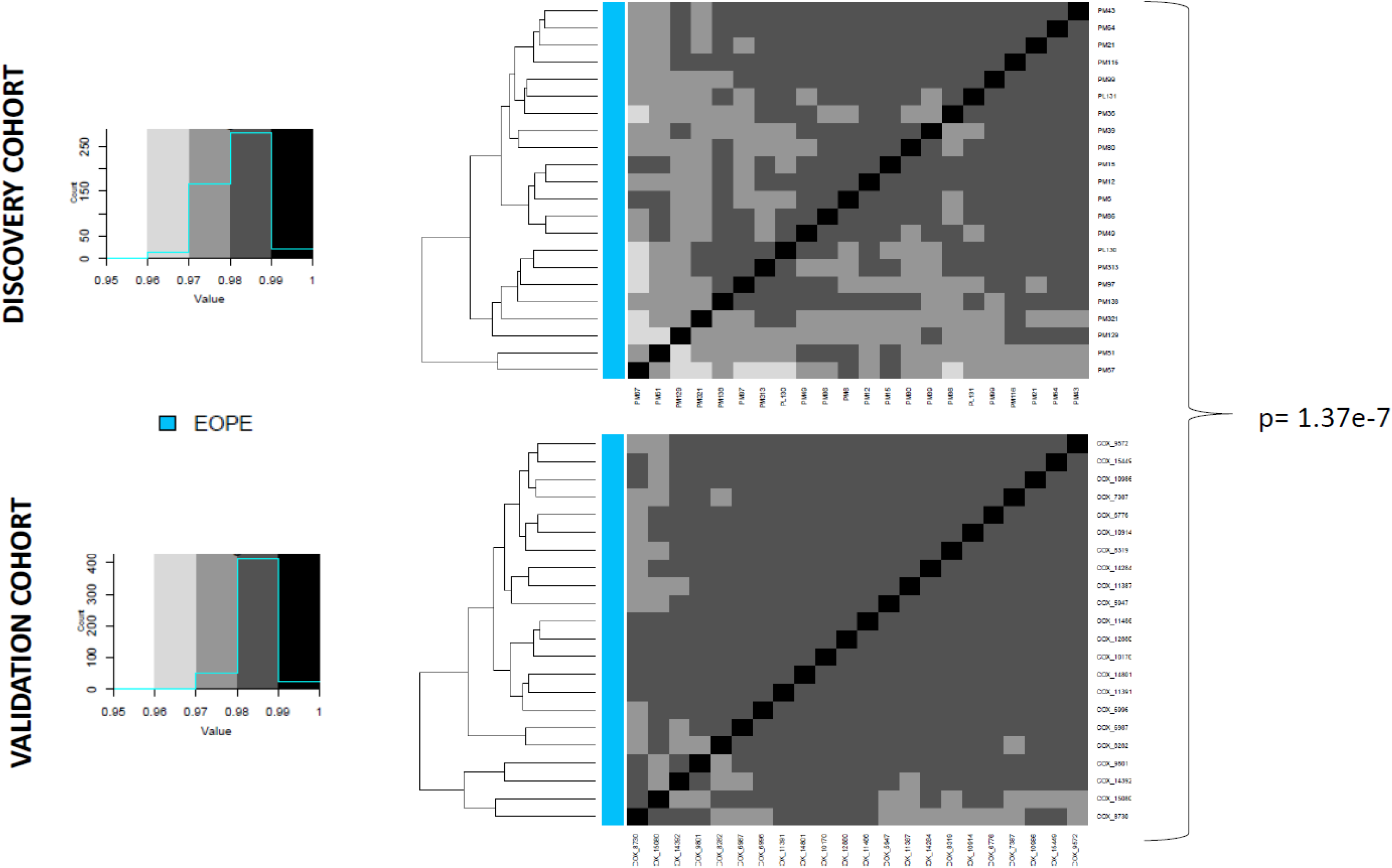
Sample-sample correlations for both the discovery and validation cohort in the EOPE samples. P-value from Student’s t-test indicates that the discovery cohort is more heterogeneous (less correlated) than the validation cohort.

**Supplementary Figure 4.**
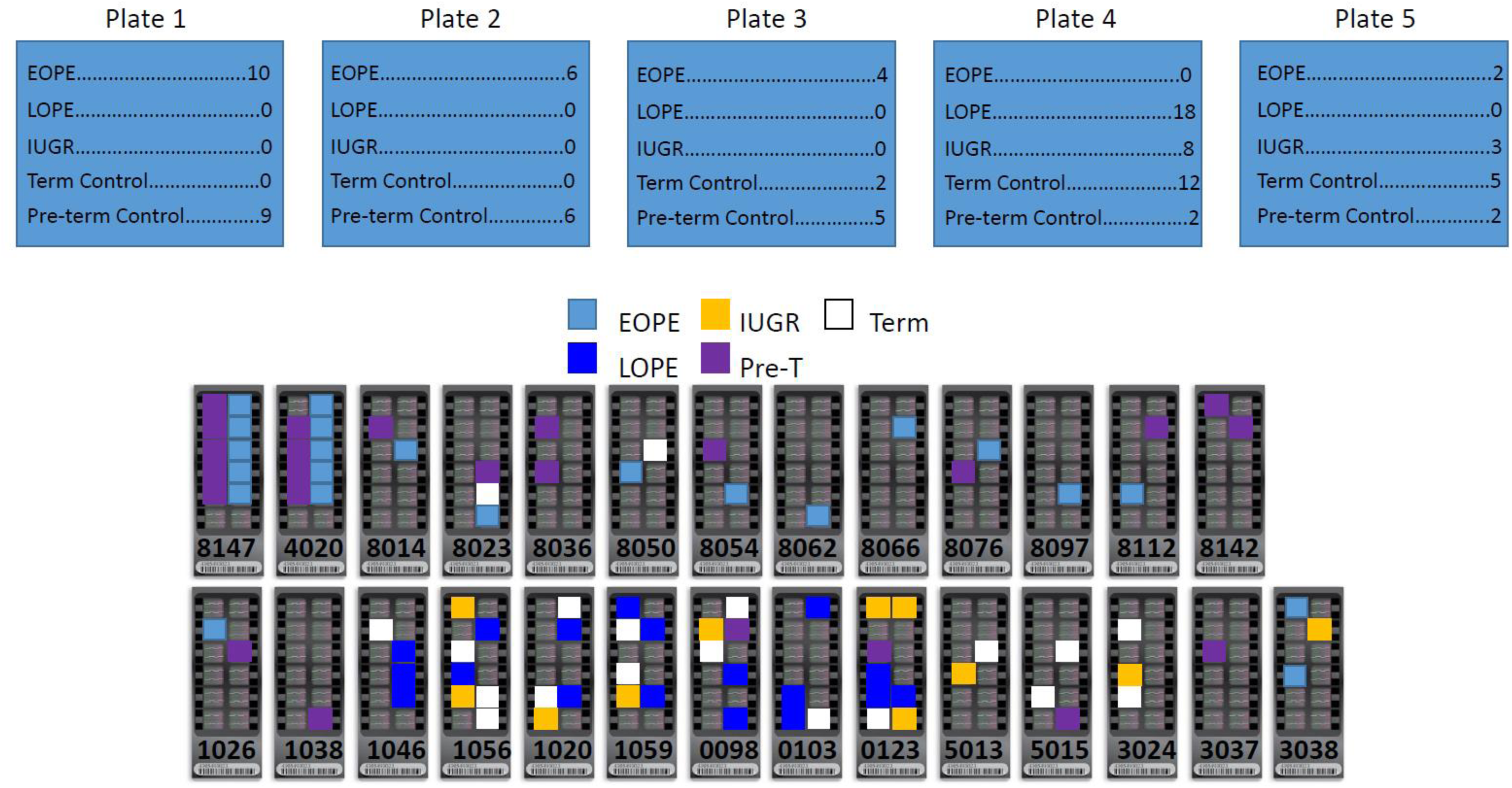
Sample placement on the 96 well plates (Plate) and the microarray chips for the discovery cohort.

**Supplementary Figure 5.**
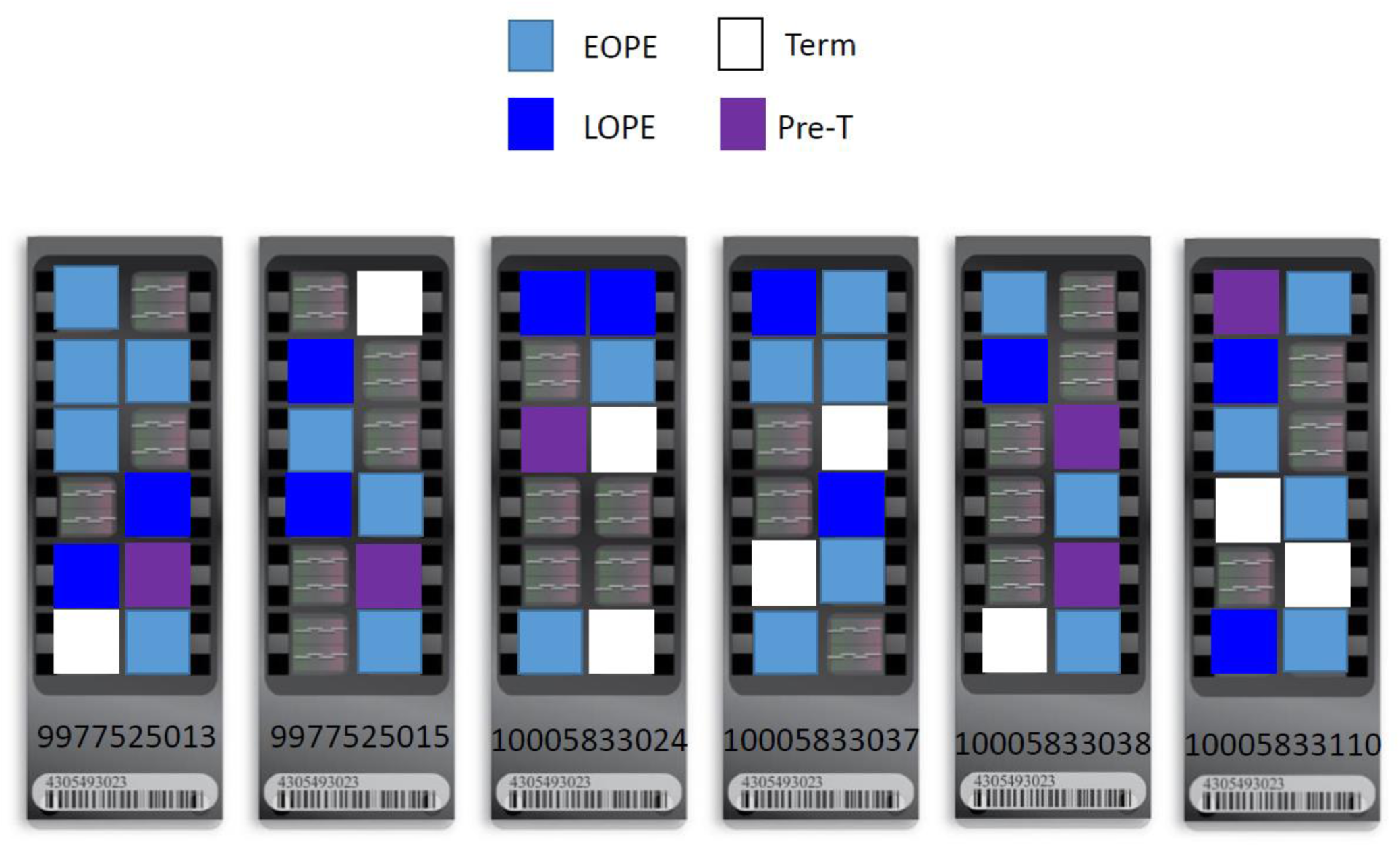
Sample placement on the 96 well plates (Plate) and the microarray chips for the validation cohort.

**Supplementary Table 1.** List of the 599 validated hits and relevant gene information

**Supplementary Table 2.** List of 244 CpG sites that are differentially methylated between methylation cluster 3 and methylation cluster 2 in both the discovery and validation cohorts.

**Supplementary Table 3.** List of 207 sites that are differentially methylation between EOPE methylation subcluster 1 and EOPE methylation subcluster 2 in both the discovery and validation cohorts.

**Supplementary Table 4.** List of probes on the 450K array that were filtered from each of the cohort.

## Abbreviations

Δβ: Delta beta
450K: Illumina infinium humanmethylation450 array DM- differentially methylated
DNAm: DNA methylation
EOPE: early-onset preeclampsia
FDR: false discovery rate
GA: gestational age
IUGR: Intrauterine growth restriction
LOPE: late-onset preeclampsia nIUGR- normotensive intrauterine growth restriction PE- Preeclampsia
PI: placental insufficiency

